# Association of zinc-finger antiviral protein (ZAP) binding to viral genomic RNA with attenuation of replication of echovirus 7

**DOI:** 10.1101/2020.05.14.097329

**Authors:** Niluka Goonawardane, Dung Nguyen, Peter Simmonds

**Affiliations:** Nuffield Department of Medicine, Peter Medawar Building for Pathogen Research, University of Oxford, Oxford OX1 3SY, UK

## Abstract

Previous studies have implicated both zinc finger antiviral protein (ZAP) and oligoadenylate synthetase 3 (OAS3)/RNASeL in the attenuation of RNA viruses with elevated CpG and UpA dinucleotides. Mechanisms and inter-relationships between these two pathways were investigated using an echovirus 7 (E7) replicon with compositionally modified sequences inserted into the 3’untranslated region. ZAP and OAS3 immunoprecipitation (IP) assays provided complementary data on dinucleotide composition effects on binding. Elevated frequencies of alternative pyrimidine/purine (CpA and UpG) and reversed (GpC and ApU) dinucleotides showed no attenuating effect nor specific binding to ZAP by IP. However, the bases 3’ and 5’ to CpG motifs influenced replication and ZAP binding; UCGU enhanced CpG-mediated attenuation and ZAP-binding while A residues shielded CpGs from ZAP recognition. Attenuating effects of elevated frequencies of UpA on replication occurred independently of CpG dinucleotides and bound non-competitively with CpG-enriched RNA consistent with a separate recognition site from CpG. Remarkably, immunoprecipitation with OAS3 antibody reproduced the specific binding to CpG- and UpA-enriched RNA sequences. However, OAS3 and ZAP were coimmunoprecipitated in both ZAP and OAS3 IP, and colocalised with E7 and stress granules (SGs) by confocal microscopy analysis of infected cells. ZAP’s association with larger cellular complexes may mediate the recruitment of OAS3/RNAseL, KHNYN and other RNA degradation pathways.

**Importance:** We have recently discovered that the OAS3/RNAseL antiviral pathway is essential for restriction of CpG- and UpA-enriched viruses, in addition to the requirement for zinc finger antiviral protein (ZAP). The current study provides evidence for the specific dinucleotide and wider recognition contexts associated with virus recognition and attenuation. It further documents the association of ZAP and OAS3 and association with stress granules and a wider protein interactome that may mediate antiviral effects in different cellular compartments. The study provides a striking re-conceptualisation of the pathways associated with this aspect of antiviral defence.

## INTRODUCTION

It is increasingly recognised that the nucleotide composition of the genomes of viruses has a profound influence of its subsequent interaction with the cell on infection. Elevated frequencies of the CpG dinucleotide in RNA virus genomes or in mRNAs expressed during replication substantially modify replication kinetics and gene expression in mammalian viruses, including enteroviruses (1) and other RNA viruses (2-5), HIV-1 (6-8) and other retroviruses (9) and hepatitis B virus (10) through their interactions with zinc finger antiviral protein (ZAP). ZAP has been shown to specifically bind to high CpG containing RNA sequences (1, 8) and activate a range of RNA degradation pathways or induce translational arrest. Amongst its many documented effects, ZAP binding may inhibit cap-dependent translation (4, 11), de-cap mRNA sequences making them susceptible to 5’-3’ degradation by the cellular Xrn1 nuclease (12, 13). ZAP additionally interacts with stress granule associated components (14-16) which may potentially sequester viral mRNAs to repress translation. Recently, it was shown that ZAP binding to CpG-enriched mRNA sequences of HIV-1 activates a cellular nuclease, KHNYN that degrades viral RNAs (6) and is required for its restriction of HIV-1 replication. In addition to KHNYN, ZAP function is additionally dependent on TRIM25, a member of the tripartite motif family of E3 ubiquitin ligases (8, 17-19).

We have recently obtained evidence that the attenuation of CpG-high mutants of the enterovirus, echovirus 7 (E7) is further dependent on oligoadenylate synthetase 3 (OAS3) and the downstream activity of RNASeL (1, 20). Reversal of attenuation was observed in both RANseL and OAS3 CRISPR k/o cell lines despite the abundant expression of ZAP in these cell lines. We have however not established how or whether CpG-high sequences might be recognised by OAS3 and then degraded by RNAseL or the nature of the interaction of ZAP with antiviral pathway. The target of OAS3 (and other OASs) are long dsRNA duplexes (21) without any substantial dependence of the RNA sequence for binding, making it unlikely that OAS directly recognises and specifically binds to high CpG RNA sequences. We speculated that OAS3 and RNASeL formed part of an alternative downstream antiviral pathway activated by ZAP, although the complete reversal of CpG-mediated attenuation of E7 in OAS3 k/o cells is seemingly at odds with the demonstrated role of KHNYN downstream of ZAP in HIV-1 restriction (6).

In addition to the growing complexity of the antiviral pathways mediated by ZAP binding, we have additionally shown that RNA sequences with elevated frequencies of the UpA dinucleotide may additionally be targeted for ZAP binding and virus restriction (1). UpA-mediated restriction in E7 was moreover similarly reversed in ZAP, OAS3 and RNAseL k/o cell lines. RNA transcripts of an E7 viral cDNA with an UpA-high insert in the coding region showed a 1000-fold greater binding to ZAP compared to WT sequences, similar to that observed for a similarly constructed CpG-high viral RNA (1). While co-crystallisation of ZAP with CpG-high ligands (22, 23) has recently provided information on the interaction of the zinc finger domains with CpG dinucleotides (and the stoichiometry of bases surrounding the motif), it is conceivable that UpA, as a similarly shaped pyrimidine (Y) – purine (R) dinucleotide might fit into the binding site. This degree of target flexibility would imply that other YpR dinucleotides (UpG and CpA) might be also recognised. An alternative possibility is that ZAP possesses an alternative recognition domain or mechanism of interaction with UpA dinucleotides.

In the current study we have made extensive use of our recently developed E7 replicon construct (20) in which compositionally modified sequences can be inserted into the 3’UTR. Because inserted sequences are not translated, any attenuation they produce cannot be mediated though previously speculated effects of CpG (and UpA) sequence modification to produce disfavoured codon pairs and reduce translation efficiency (24-28). The system additionally provides much greater freedom to manipulate sequence composition of the inserts since they do not need to retain the amino acid coding of the native sequence. This restriction greatly limits the mutagenesis possible in currently described virus models. The replicon system allowed us to investigate effects of modification of other dinucleotide frequencies, specifically the alternative YpR dinucleotides, reversed dinucleotides (GpC and ApU) and effects of altering the upstream and downstream bases (context) of CpG dinucleotides. The latter was motivated by the possibility of a broader recognition motif for ZAP binding. Effects of sequence modifications on replicon replication were compared with binding affinities of the replicon RNAs to ZAP and OAS3 in an *in vitro* binding assay. This assay enabled further studies of the site specificity of CpG and UpA binding to ZAP and the existence of shared or separate binding sites. Finally, through the use of cell lysates from ZAP and OAS3 k/o cell lines we were able to investigate the inter-dependence of ZAP and OAS3 on RNA binding and their potential cellular interactions.

## RESULTS

### Alternative dinucleotides in virus attenuation

Replication kinetics of E7 replicons are largely unaffected by genome length (20), enabling effects of relatively large insertions of compositionally modified sequence on attenuation to be determined. The current study made extensive use of the replicon construct with a 1242 nucleotide sequence inserted into the 3’UTR (ncR1; Fig. 1A). Using this model, we investigated whether elevated frequencies of CpA and UpG dinucleotides attenuated replication similarly to the alternative YpR dinucleotides, UpA and CpG. Sequences with maximised frequencies of each were synthesised (Table 1); these contained approximately 3-fold more CpA or UpG dinucleotides than the WT sequence, the maximum possible while keeping mononucleotide frequencies and those of UpA and CpG constant.

**TABLE 1.** Composition of 3’UTR and coding region mutants of E7^1^

| Region<sup>2</sup> | Mutant | Coding<sup>3</sup> | Length | f_G+C | Dinucleotide totals<sup>4</sup> |  |  |  |  |  |
| --- | --- | --- | --- | --- | --- | --- | --- | --- | --- | --- |
|  |  |  |  |  | CpG | UpA | ApU | GpC | CpA | UpG |
| Standard composition mutants |  |  |  |  |  |  |  |  |  |  |
| 3'UTR | WT | Yes | 1233 | 47.6% | 51 | 62 | 85 | 70 | 130 | 92 |
| 3'UTR | CpG-High | Yes | 1233 | 56.5% | 180 | 52 | 64 | 116 | 78 | 51 |
| 3'UTR | UpA-High | Yes | 1233 | 40.9% | 39 | 171 | 114 | 53 | 76 | 69 |
| Reversed dinucleotides |  |  |  |  |  |  |  |  |  |  |
| 3'UTR | ApU-Max | No | 1233 | 47.6% | 51 | 62 | 205 | 70 | 122 | 97 |
| 3'UTR | GpC-Max | No | 1233 | 47.6% | 51 | 62 | 85 | 180 | 118 | 102 |
| Alternative pyrimidine purine dinucleotides |  |  |  |  |  |  |  |  |  |  |
| 3'UTR | CpA-Max | No | 1233 | 47.6% | 51 | 62 | 114 | 96 | 191 | 91 |
| 3'UTR | UpG-Max | No | 1233 | 47.6% | 51 | 62 | 104 | 91 | 130 | 171 |
| Altered CpG 5' & 3' base contexts |  |  |  |  |  |  |  |  |  |  |
| 3'UTR | ACGA_WT | No | 1233 | 47.6% | 51 | 62 | 76 | 47 | 87 | 91 |
| 3'UTR | CCGC_WT | No | 1233 | 47.6% | 51 | 62 | 85 | 96 | 89 | 90 |
| 3'UTR | GCGG_WT | No | 1233 | 47.6% | 51 | 62 | 83 | 83 | 109 | 64 |
| 3'UTR | UCGU_WT | No | 1233 | 47.6% | 51 | 62 | 73 | 47 | 105 | 70 |
| 3'UTR | ACGA_CpG-H | No | 1233 | 47.6% | 102 | 62 | 80 | 44 | 72 | 88 |
| 3'UTR | CCGC_CpG-H | No | 1233 | 48.1% | 102 | 62 | 97 | 117 | 52 | 87 |
| 3'UTR | GCGG_CpG-H | No | 1233 | 46.9% | 102 | 62 | 102 | 119 | 103 | 37 |
| 3'UTR | UCGU_CpG-H | No | 1233 | 47.6% | 102 | 62 | 66 | 28 | 102 | 46 |
| 3'UTR | xCGx | No | 1233 | 54.4% | 102 | 62 | 65 | 89 | 56 | 95 |
| Virus coding region mutants |  |  |  |  |  |  |  |  |  |  |
| R1 | CpGH_UpAL | Yes | 1233 | 56.2% | 179 | 19 | 62 | 102 | 95 | 53 |
| R1 | CpGL_UpAH | Yes | 1233 | 41.0% | 0 | 170 | 114 | 52 | 82 | 84 |

<sup>1</sup>A full listing of the composition of these sequences is provided in Table S3; Suppl. data<sup>2</sup>Genome region mutated<sup>3</sup>Retention of native coding<sup>4</sup>Total of dinucleotides or base frequencies of mutated sequences. Figures in grey boxes represent deliberately altered values.

**FIGURE 1.**
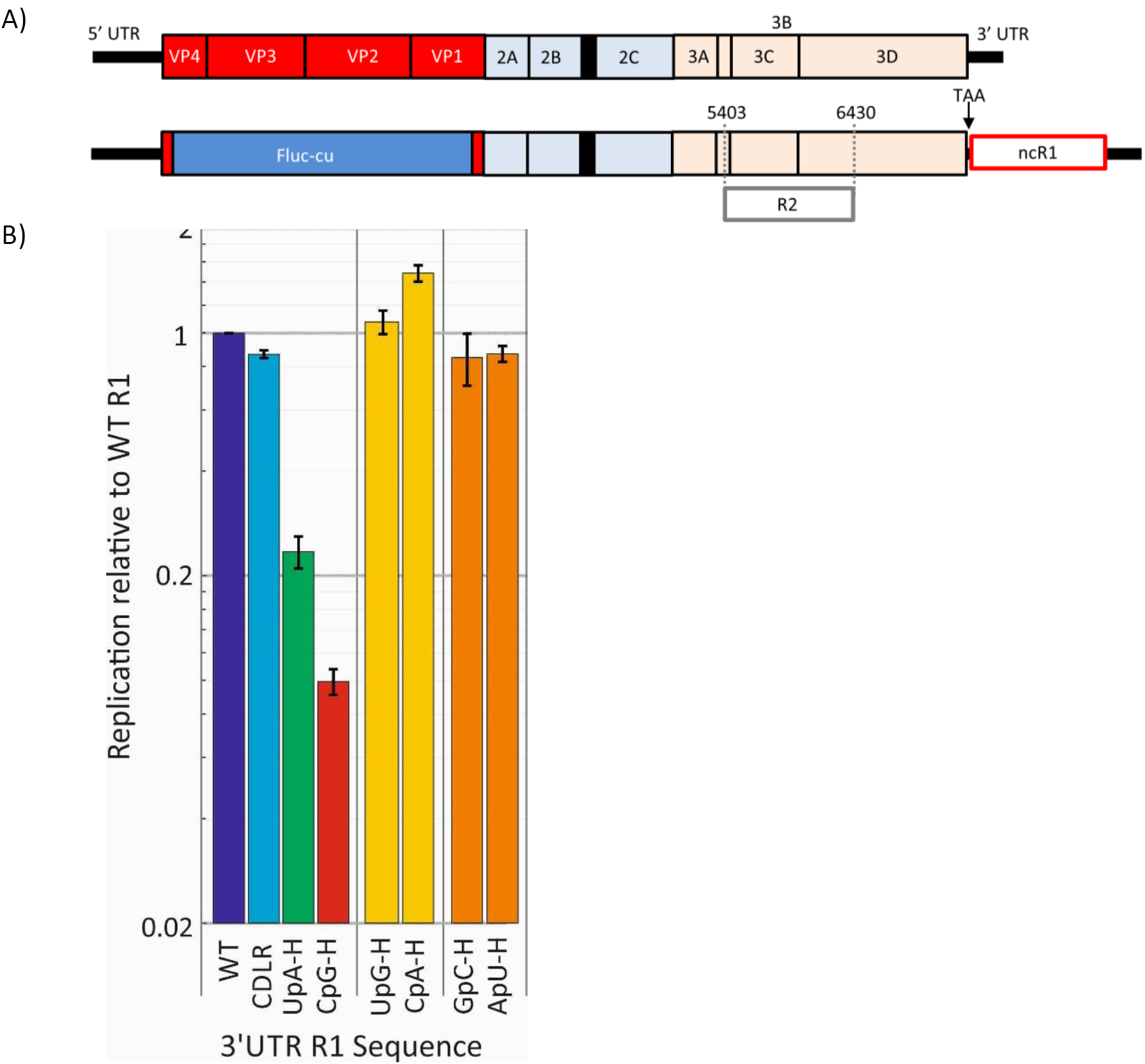
REPLICON MODEL USED TO INVESTIGATE EFFECTS OF DINUCLEOTIDE FREQUENCY MODIFICATIONS OF R1 SEQUENCES IN THE 3’UTR ON E7 REPLICON REPLICATION. A) Schematic representation of genome organization of E7 virus (top) and the replicon with non-coding synthetic region R1 (ncR1) inserted into the 3’UTR. (B) Replication of the replicons in which the ncR1 region has been modified to increase frequencies of CpG, UpA, the other pyrimidine / purine dinucleotides (UpG and CpA) and reversed dinucleotides (GpG and ApU). Replication was measured by luciferase expression at 6 hours p.t. and expressed as a ratio of replicon luminescence from a replicon with the E7 WT ncR1 sequence (normalised to 1.0). Bar heights represent the mean of two biological replicates (each the mean of three technical replicates). Error bars show standard deviations (SDs).

We additionally investigated whether the attenuation created by CpG and UpA-high mutants originates from the resulting increased frequency of self-complementary base pairs in viral genomic RNA or mRNAs. Two further additional mutants were generated with sequences containing maximised frequencies of reversed dinucleotides (GpC and ApU). These retained native frequencies of CpG and UpA but their altered composition would have a similar effect on RNA structure formation in the virus (or replicon) genome. Bioinformatically, increasing the frequencies of GpC and ApU dinucleotides substantially increased the minimum free energy (MFEs) on folding predicted by RNAFold from −60.9 KCal/mol of the WT sequences to −72.4 and −78.5 KCal/mol of the ApU-H and GpC-H mutants respectively (Table S3; Suppl. Data). These were comparable to values of −76.4 and −69.9 KCal/mol of the equivalently mutated UpA-H and CpG-H sequences.

RNA from both sets of mutants was transcribed from linearised replicon cDNA clones along with previously described WT, CpG-H and UpA-H mutants as controls (20). RNA was transfected into A549 cells and replication was monitored by luciferase expression at different time point post transfection (p.t.). At 6 hours p.t., there was an approximate 20- and 4-fold difference in luciferase expression between the WT replicon and those with the CpG-high and UpA-high inserted sequences (Fig. 1B). At this time point, however, replication of all 4 newly constructed mutants (ApU-Max, GpC-Max, CpA-Max and UpG-Max) was comparable to the WT E7 replicon. The template sequences created to maximise frequencies of the four dinucleotides were used to produce CpG and UpA-high mutants that were otherwise equivalent in sequence composition (Fig. S1 and Table S3; Suppl. Data). In contrast to replicons with ApU-Max, GpC-Max, CpA-Max and UpG-Max inserts, the replication of both sets of CpG-Max and UpA-Max sequences attenuated to a similar degree to those of the original, coding preserved CpG-H and UpA-H mutants (Fig. S1; Suppl. Data).

Together, these findings provide no evidence for recognition and downstream potential ZAP-mediated restriction of replication of the CpA and UpG mutants. Furthermore, the suspected effects of increased frequencies of GpC and ApU dinucleotides on RNA configuration has no effect on their replication and this mechanism is unlikely to account for the attenuation of the CpG- and UpA-high mutants.

### Effects of 5’ and 3’ base contexts on CpG-mediated attenuation

To investigate whether upstream and downstream contexts influenced CpG-mediated attenuation of the E7 replicon, two sets of modified synthetic sequences with A, C, G or U bases surrounding each CpG dinucleotide were created. The first set contained the same number (n=51) and positions (and therefore spacings) of CpG dinucleotides as found in the native R1 sequence; a second set was generated in which the number of CpG dinucleotides was doubled, each in their specific 5’ and 3’ contexts.

Transfection of RNA transcripts from these constructs revealed marked differences in replication kinetics that was strongly influenced by CpG sequence context (Fig. 2A). For both sets of mutants (51 and 102 CpG dinucleotides), those with CpGs surrounded with U residues were substantially more attenuated that those in an A context, with C and G contexts being intermediate. Remarkably, surrounding CpG with U residues was sufficient to profoundly restrict the replication of the mutant with WT numbers of CpGs (n=51). Conversely, surrounding CpGs with A residues prevented any attenuation even of the CpG high mutant (n=102). A similar but even more extreme context effect on attenuation was observed in RD cells (Fig. S2; Suppl. Data)

**FIGURE 2.**
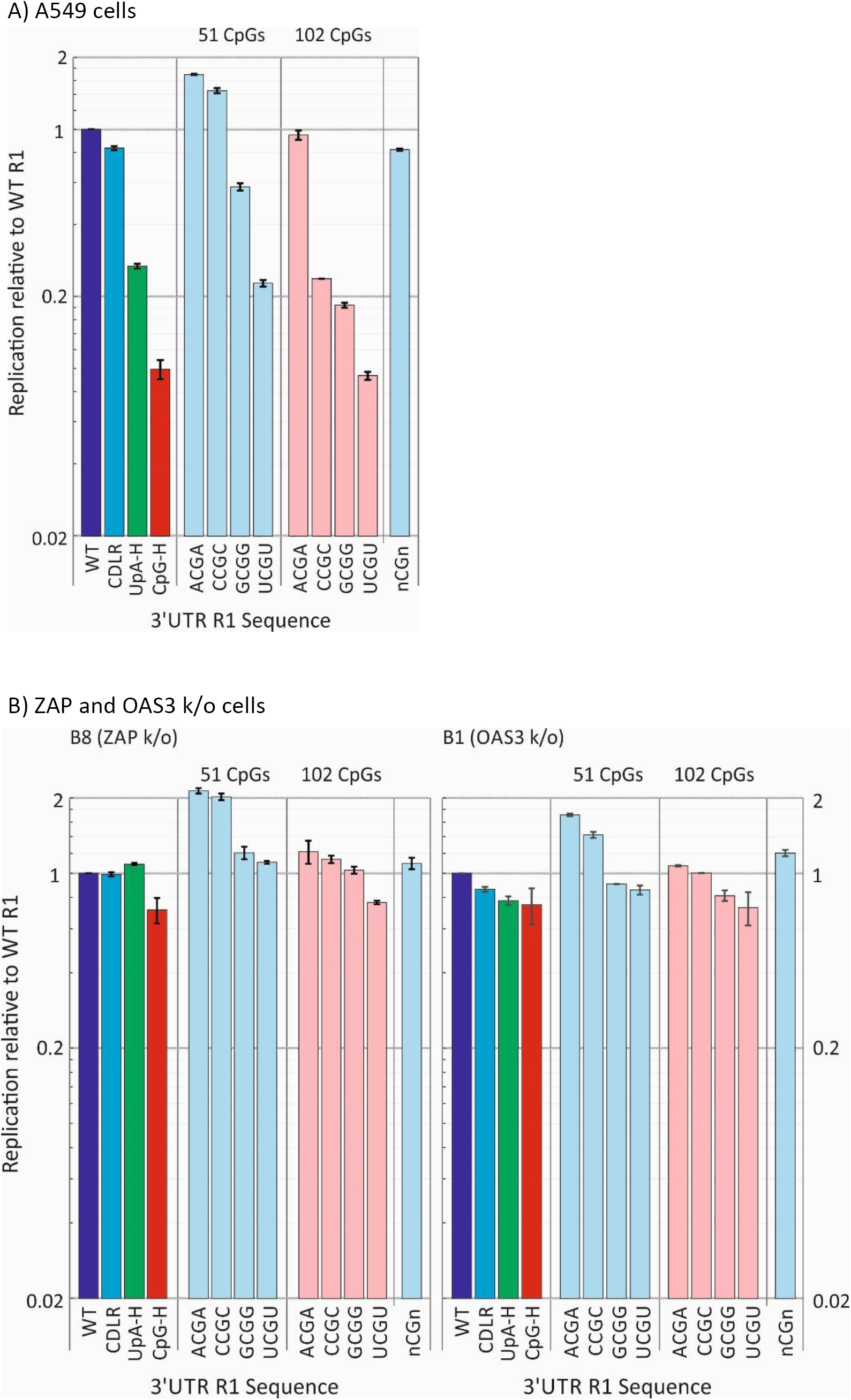
EFFECT OF 5’ AND 3’ BASE CONTEXTS OF CpG ON REPLICATION. Replication of replicons with R1 sequences containing WT or increased numbers of CpG dinucleotides (n=51 and n=102 respectively) with specified 5’ and 3’ bases (contexts) - A, C, G, T, or n (randomised). Replication at 6 hours was normalised to WT replication (WT = 1.0). CpG-H and UpA-h mutants that previously showed attenuated replication were included as controls. Bars represent the mean of two biological replicates, each with three technical replicates, error bars show SDs. (A) Replication in A549 cells. (B) The equivalent experiment performed in ZAP and OAS3 k/o cells.

To investigate the roles of ZAP and OAS3 in the varied attenuation phenotypes of these mutants, the transfections were repeated in B8 (ZAP k/o) and B1 (OAS3 k/o) cells (Fig. 2B). For both cell lines, replication differences between mutants were eliminated providing an indication that the contexts of the CpG dinucleotides influenced their recognition by ZAP (and OAS3) and that the differences in replication did not arise through an alternative restriction mechanism.

The effects of sequence context on CpG-mediated attenuation were similarly apparent when replication was assayed by immunofluorescent detection of dsRNA in infected cells (Fig. 3). The replicon with a WT sequence inserted into the 3’UTR showed high levels of accumulation of dsRNA and induction of ZAP expression at 4 hours p.t.. Transfection of A549 cells with the CpG-H and UpA-H mutants induced far less dsRNA, consistent with their attenuation of replication in other assays (Fig. 1B). The different degrees of attenuation of the context mutants (ACGA, CCGC, GCGG, UCGU) in the replication assay (Fig. 2) was similarly reflected in amounts of dsRNA detected by IF (Figs 3B, 3C). Quantitation of dsRNA staining intensity by Airyscan ranked dsRNA detection comparably to replication (Fig. 3D).

**FIGURE 3.**
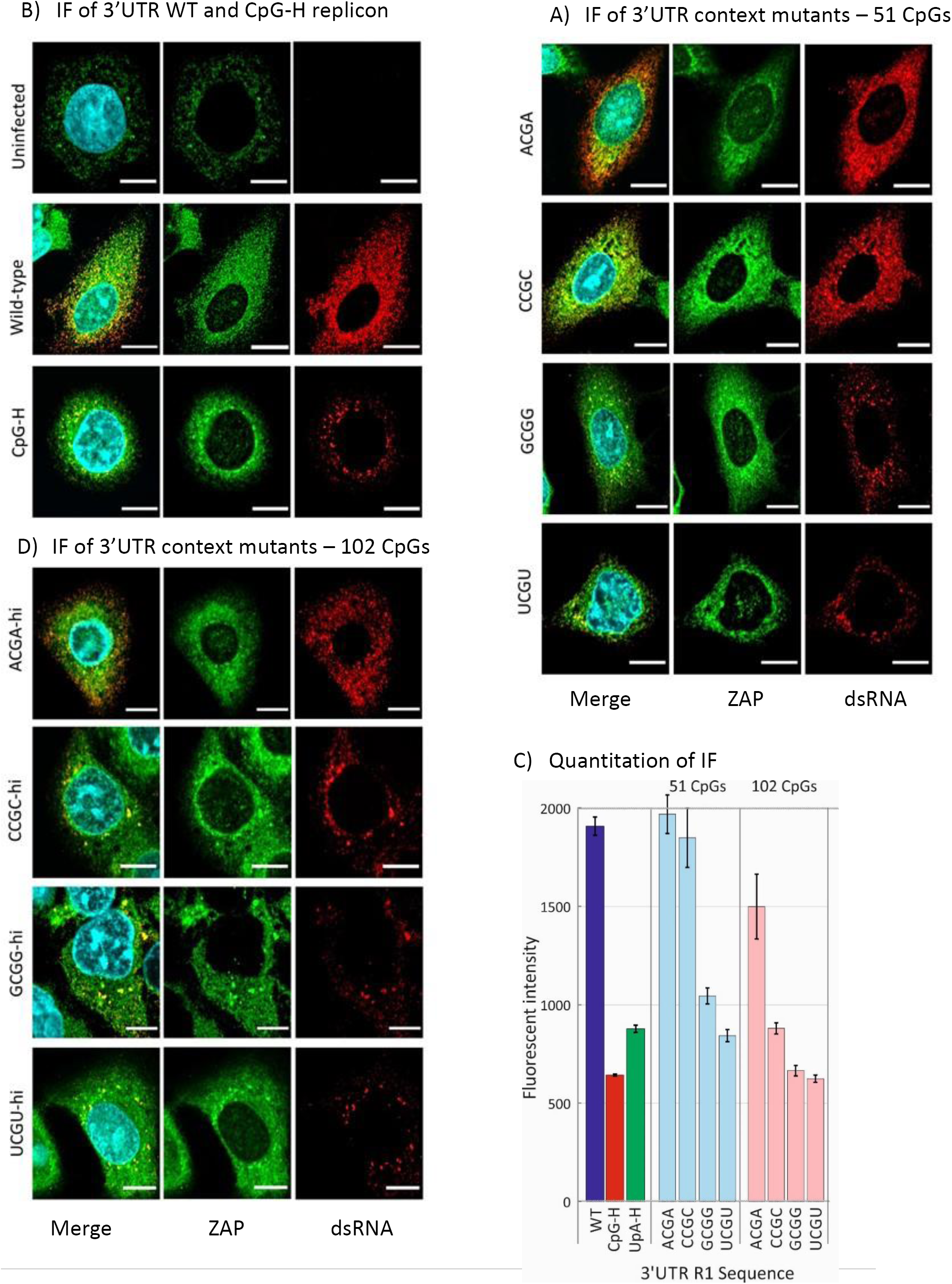
IF DETECTION OF dsRNA (REPLICATING E7) AND ZAP EXPRESSION IN TRANFECTED A549 CELLS. (A) Uninfected A549 cells co-stained for ZAP by specific antibody and Alexa Fluor 488-coupled secondary antibody and for E7 RNA by J2 (assay specificity control, not detected). WT or CpG-H (assay positive control) transfected cells co-stained for ZAP (green) and for dsRNA detected (red hot) by J2 antibodies in cells at 6 h post transfection. Nuclear DNA was stained by DAPI (blue). 3’UTR mutant sequences with either (B) WT level (n=51) or (C) elevated (n= 102) CpG frequencies. Scale bars: 10 μm. (D) Quantitation of fluorescent intensity of infected cells by Airyscan post-acquisition analysis in representative fields of cell monolayer transfected with E7 WT and compositionally modified mutants of E7. Bar heights show the mean of two biological replicates; error bars show ± 1 SD.

### Influence of dinucleotide composition on RNA binding to ZAP

We applied a previously described *in vivo* binding assay for ZAP in which viral RNA sequences transfected into cells were extracted into a cell lysate and the fraction bound to ZAP estimated through immune capture with an anti-ZAP antibody (Fig. 4). The CpG-H and UpA-H control showed enhanced binding to ZAP compared to the WT replicon, consistent with previous observations (1). As observed previously, the UpA-H sequence bound more avidly than CpG-H. Replicon mutants with the alternative YpR (CpA and UpG) or reversed (GpC and ApU) dinucleotides showed binding similar to that of the WT replicon, consistent with their lack of attenuation of replication in A549 cells (Fig. 2A). However, the binding or replicon RNA with the CpG dinucleotides placed in different contexts showed markedly different binding, greatest for the UCGU motif and least with ACGA. The findings recapitulate the previous observations of their relative attenuation, including the finding that the mutant with elevated frequencies of CpG motifs surrounded by A residues bound similarly to the WT control, while those with the UCGU motif bind over ten-fold more strongly to ZAP even with WT number of CpG dinucleotides. From these observations, effects of sequence context on attenuation may primarily originate from effects on ZAP binding.

**FIGURE 4.**
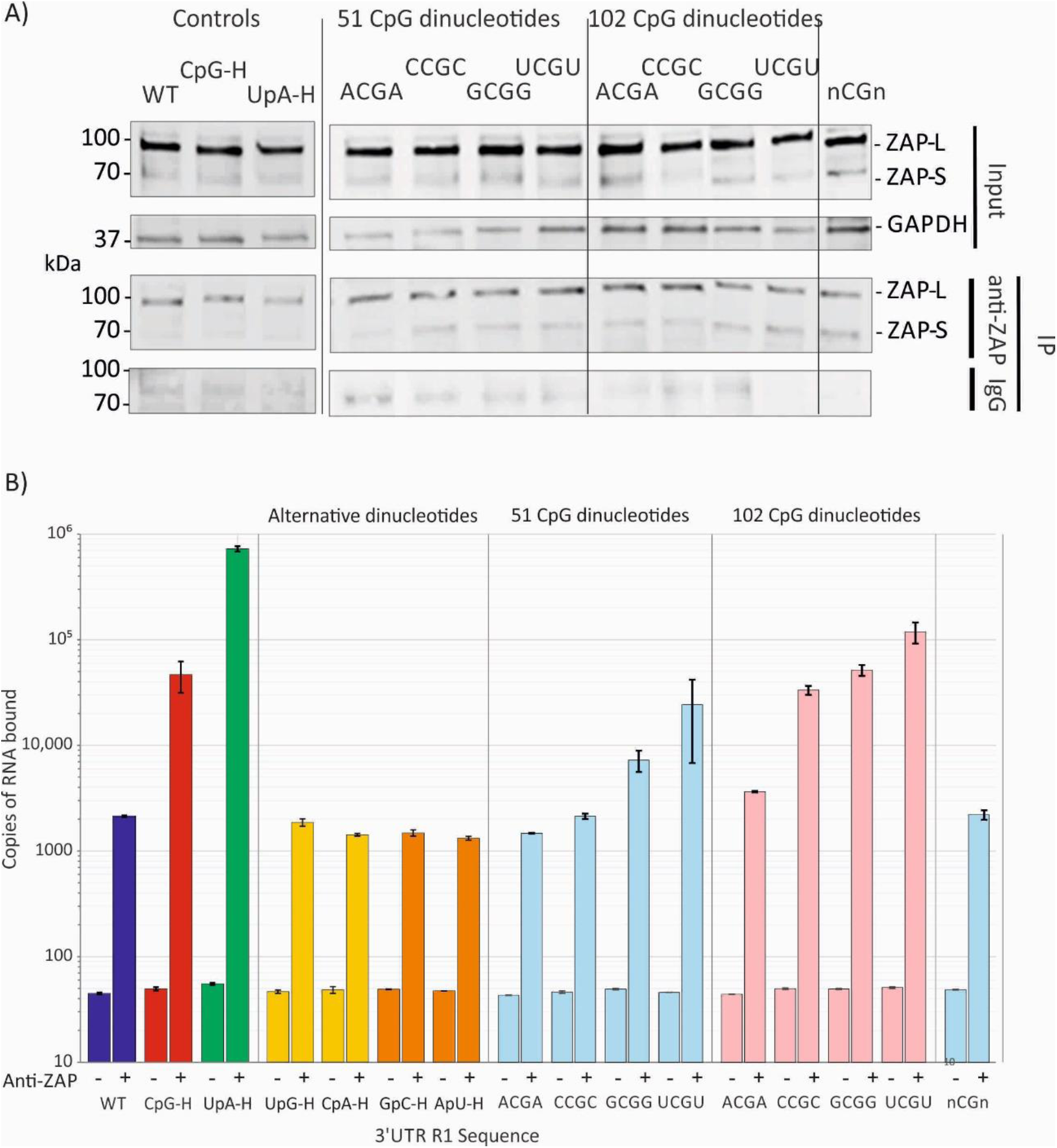
BINDING OF REPLICONS TO ZAP AND OAS3 BY COLUMN IMMUNOPRECIPITATION. Binding of ZAP to WT, UpA-H and CpG-H controls or 3’UTR mutant RNA transcripts. (A) WB of input long (L) and short (S) isoforms of ZAP (∼100 kDa and ∼70 kDa respectively) and GAPDH in uninfected cell lysates detected by immunostaining with anti-ZAP polyclonal antibody or GAPDH polyclonal antibody. Cell lysates from replicon transfected cells were incubated with anti-ZAP Ab or control antibody (IgG) and immobilised to columns with anti-rabbit IgG. The lower panel shows a WB for immunoprecipitated ZAP. (B) Quantitation of E7 viral RNA transcripts with WT, CpG-H and UpA-H or 3UTR mutant sequences by qPCR in immunoprecipitated ZAP (right column) or mock precipitated control (left column). RNA was quantified by qPCR using conserved primers and probe from the 5’UTR (Table S2; Suppl. Data). Bar heights and error bars represent means and SDs of two biological replicates.

To more directly evaluate the propensity of ZAP to bind to RNA sequences enriched for CpG and UpA, we devised an *ex vivo* immunoprecipitation assay format in which uninfected cell lysates were incubated in a reaction mix containing anti-ZAP antibodies and RNA transcripts of the R1 region. The ZAP/RNA complexes were subsequently immobilised, stringently washed and bound RNA quantified using primers matching the insert sequences (Fig. 5). ZAP binding to R1 RNA sequences showed a similar greater (X100-fold) affinity for CpG- and UpA-enriched sequences as observed for E7 containing these sequences (Fig. 4). This assay allowed us to investigate whether the lower but above background observed level of ZAP binding to WT E7 sequences (compared to the no antibody control) resulted from the presence of a lower but non-zero number of CpG and UpA dinucleotides in the sequence (n=51 and 49 respectively). However, binding of ZAP was comparable to the CpG-zero, UpA-low and combined CpG-zero/UpA-low sequences (Fig. 5, right panel). These findings suggest a level of sequence independent binding of ZAP to RNA sequences.

**FIGURE 5.**
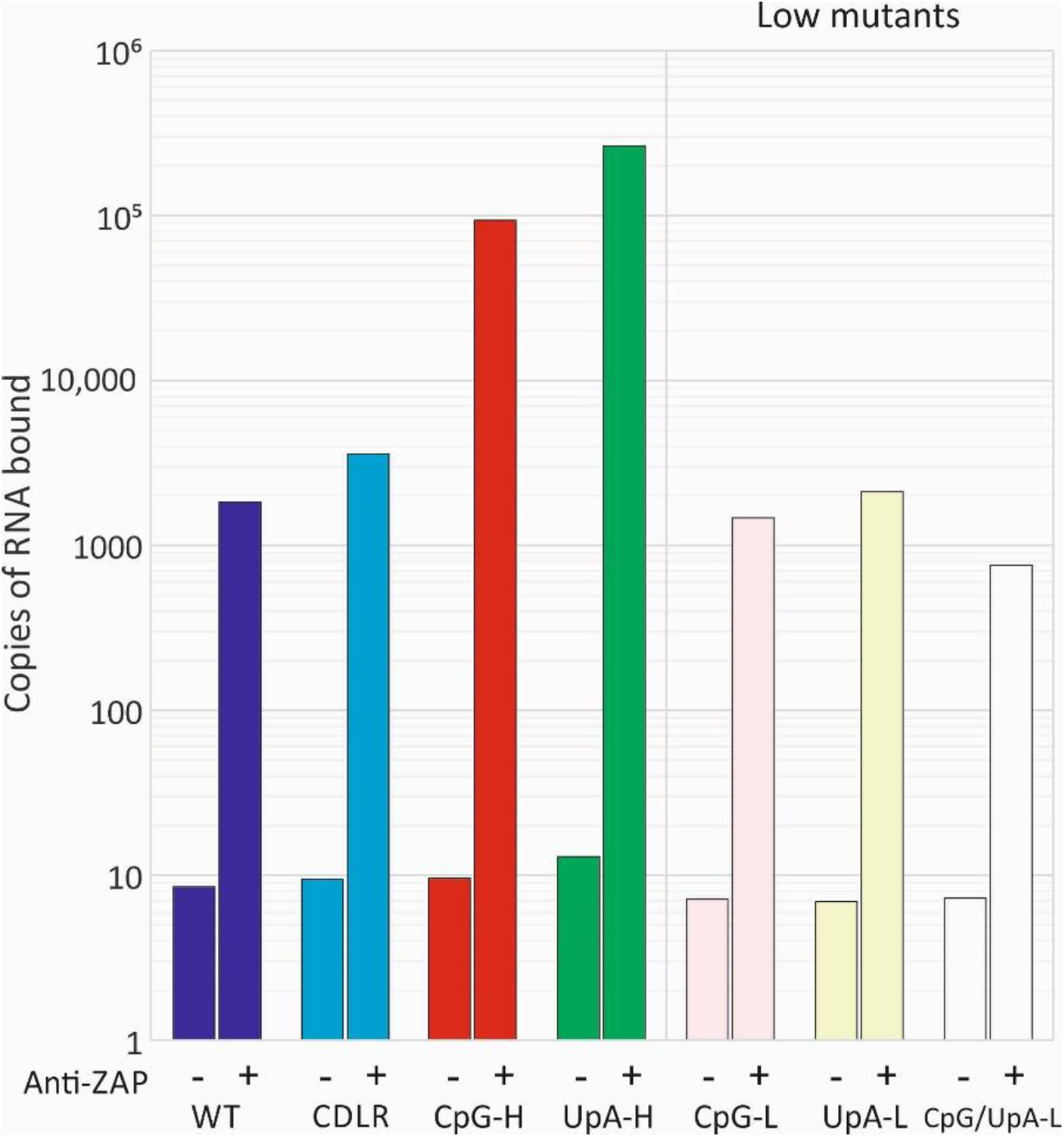
BINDING OF ZAP TO R1 RNA SEQUENCES OF DIFFERENT COMPOSITION *EX VIVO*. Binding of ZAP in IFN-β pre-stimulated cell lysates to RNA of different compositions *ex vivo*. **+**: ZAP immunoprecipitated with anti-ZAP antibody; **-**: mock precipitated control. RNA was quantified by qPCR using R1 sequence-specific primers for the different mutants (UpA-H, CpG-H, UpA-L and combined WT, CpG-L and CpG/UpA-L; Table S2; Suppl. Data).

### Characterisation of ZAP binding to UpA-enriched RNA sequences

In this (Fig. 2A) and in previous studies (1), both viruses and replicons with increased UpA numbers either in coding regions or the 3’UTR showed ZAP-dependent attenuation. UpA-H sequences within a viral RNA transcript or a replicon (Fig. 4B) additionally showed greater binding to ZAP that the WT constructs, in both cases with greater affinity than a sequence comparably enriched for CpG. Since enrichment for CpA and UpG did not attenuate E7 replicons (Fig. 1B), it seems unlikely that ZAP binding to UpA arose simply through a reduced selectivity of the dinucleotide binding site in ZAP for YpR (*ie.* any pyrimidine followed by any purine). To investigate whether CpG and UpA binding occurred through separate binding sites, we performed a competition assay in which CpG binding was assayed in the presence of differing concentrations of a UpA enriched sequence and *vice versa* (Fig. 6).

**FIGURE 6.**
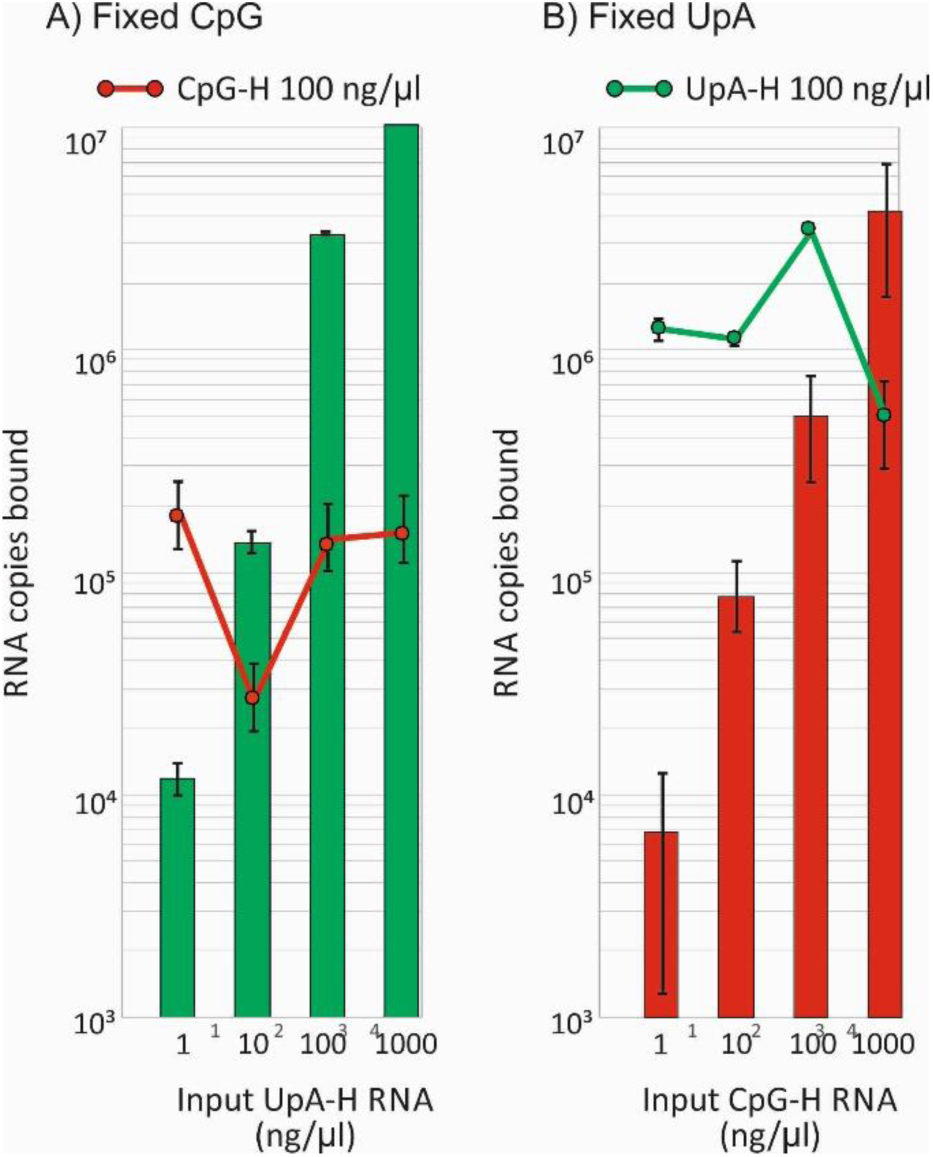
COMPETITION ASSAY FOR CpG AND UpA BINDING TO ZAP. Co-incubation of replicon RNA with fixed concentrations (100 µg/µl) of either (A) CpG-H or (B) UpA-H replicons in the presence of varying concentration of the heterologous replicon. RNA binding was quantified by CpG-bor UpA-specific PCR to enable transcripts to be differentiated (Fig. S3; and Table S2; Suppl. Data). Bar heights show the mean of two biological repeats; error bars show SDs.

The binding of the CpG-H replicon was unaffected by co-incubation with differing concentrations of the UpA-H replicon, even if in 10-fold molar excess. Similarly, the binding of the UpA-H replicon was unaffected by co-incubation with the CpG-H replicon over the same range of relative concentrations. These findings are most consistent with the existence of separate binding sites for CpG and UpA dinucleotides in ZAP or associated, co-immunoprecipitated proteins.

Previous investigations of ZAP and UpA-mediated effects on E7 and other virus replication have not considered whether both efficient binding and ZAP-mediated attenuation of replication requires the co-presence of both CpG and UpA dinucleotides in the same RNA sequence. The observation for the potential existence of separate recognition sites does not rule out this possibility. To investigate this we constructed a further pair of replicon mutants in which the insert 3’UTR sequence was constructed to contain either zero CpG dinucleotides and maximised numbers of UpAs – and *vice versa* (Fig. 7). The replication of the CpG-H / UpA-L mutant was substantially attenuated despite almost complete absence of UpA dinucleotides. Conversely, the UpA-H / CpG-zero construct was similarly attenuated compared to the control UpA-H mutant that possessed close to WT levels of CpG. Both mutants showed slightly less attenuation than the original mutants although this was proportional to the lesser degree of CpG and UpA elevation possible while retaining mononucleotide frequencies and coding in these mutants. Overall these findings show that either dinucleotide can independently attenuate replication.

**FIGURE 7.**
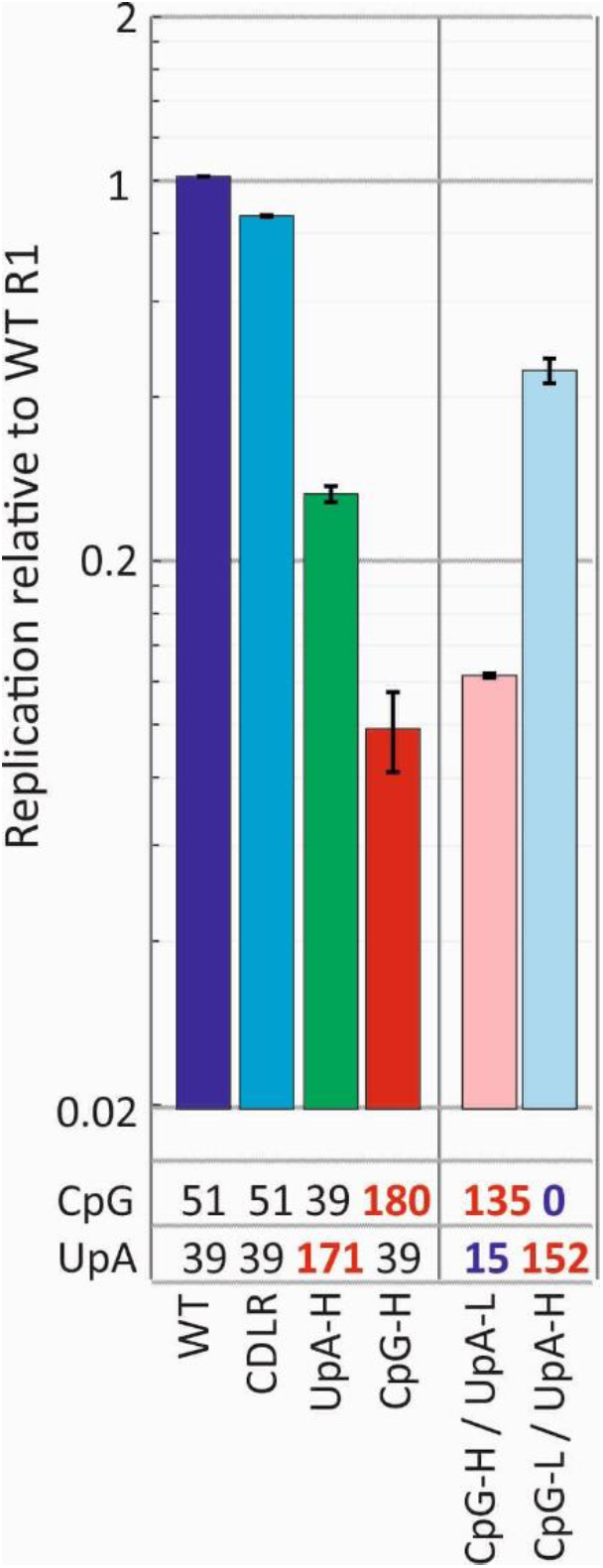
INDEPENDENCE OF CpG AND UpA DINUCLEOTIDES ON REPLICON ATTENUATION. Replication of mutants of E7 replicons with minimised and maximised CpG and UpA dinucleotides. The totals of each dinucleotide in each 3’UTR sequence are indicated under the graph – bold numbers indicate total that has been deliberately maximised and minimised (for these mutants while retaining protein coding). Bar heights show the mean of two biological repeats; error bars show SDs.

### Binding of OAS3 to compositionally modified RNA

We previously observed that expression of OAS3 was required or ZAP-mediated restriction of high-CpG and –UpA mutants of E7 viruses and replicons. The nature of the interaction between ZAP and OAS3, whether direct or indirect, is unknown although as we reviewed, it appeared unlikely structurally that the RNA binding site in OAS3 would show any affinity for the single-stranded genomic or transcript sequences of E7 we have studied to date (1). To investigate this experimentally, we modified the *ex vivo* IP assay to pull out OAS3 using an anti-OAS3 specific antibody and investigated its binding to E7 R1 RNA sequences of different compositions (Fig. 8A). Surprisingly, OAS3 demonstrated a comparable propensity to specifically bind to CpG-H and UpA-H RNA sequences to that of the ZAP IP assay (Fig. 5).

**FIGURE 8.**
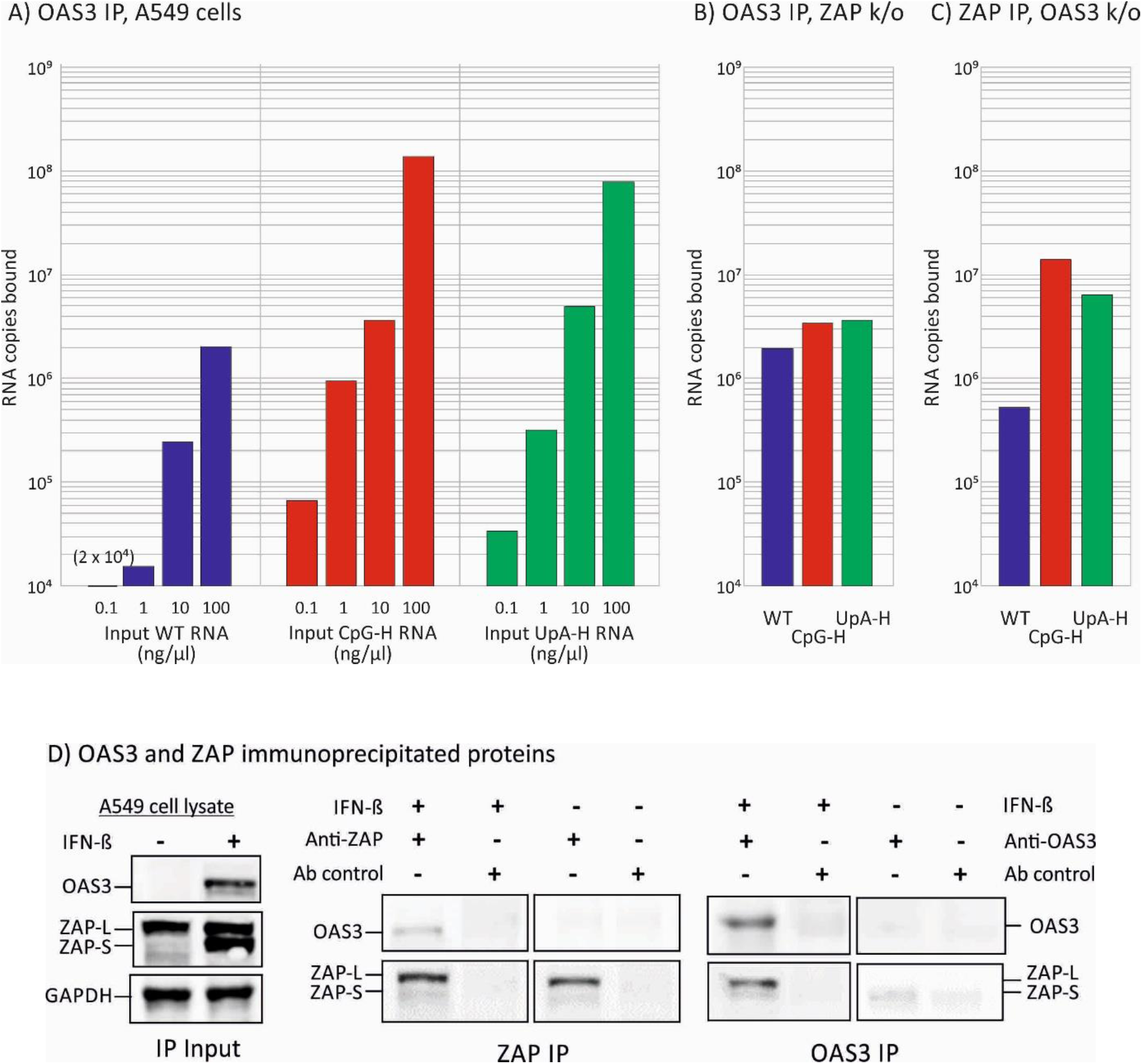
E7 REPLICON BINDING IN OAS3 IP. A) Pull-down of different input quantities of E7 R1 insert RNA sequences by OAS3; binding quantified by qPCR. (B, C) Pull-down of 100 ng input WT, CpG-H and UpA-H RNA transcripts in ZAP k/o cells and in OAS3 k/o cells respectively. (D) Western blot detection of ZAP and OAS3 by specific antibodies in cell lysates with and without IFN-β pre-stimulation (left panel) and after immunoprecipitation by ZAP and OAS3 antibodies (centre and right panels). In both IPs, ZAP and OAS3 (after IFN- β induction) co-precipitated indicating their physical interaction.

However, the ZAP-dependence of the phenomenon was demonstrated by the absence of dinucleotide-specific binding to R1 RNA in an OAS3 IP in the ZAP k/o (B8) cell line (Fig. 8B), where binding affinities to the CpG-H and UpA-H transcripts were comparable to that of the WT sequence. Contrastingly, a degree of sequence-specificity was observed in the complementary experiment – ZAP-IP of OAS3 k/o cells. These results suggest that RNA binding in the OAS3 IP is mediated through the co-precipitation of ZAP (Fig. 8C). To analyse this potential cellular interaction further, we probed the OAS3 immune precipitate for ZAP, and vice versa (Fig. 8D). Cell lysates from A549 cells showed constitutive expression of the long (L) isoform of ZAP while the expression of the short (S) isoform and of OAS3 was minimal or undetectable. Both were potently induced after pre-treatment with IFN-β (Fig. 8D, left panel). In the absence of E7 replicon RNA, subsequent immunoprecipitation of the cell lysate with anti-ZAP antibody also immunoprecipitated OAS3 after IFN stimulation (middle panel). Similarly, IP with anti-OAS3 antibody co-immunoprecipitated both isoforms of ZAP (right hand panel). Their demonstrated physical association in the *ex vivo* conditions used in the IP assay indicates that apparent binding of CpG-UpA-high RNA in the OAS IP assay (Fig. 8A) may have been mediated by ZAP or other proteins co-immunoprecipitated in the assay (See Discussion).

Further evidence for the close association of ZAP and OAS3 in infected cells was provided by confocal microscopy of E7 transfected cells and immunostaining for ZAP, OAS3, E7 and G3BP (Fig. 9). Virus replication was localised by detection of cytoplasmic dsRNA by specific antibody (J2; SCICONS) in preference to our previous detection by antibodies to a viral capsid protein; the locations of assembled virions may not coincide precisely with sites of virus replication. WT and CpG-H transfected replicons induced different levels of replication at 6 hours (Fig. 9A, 9B) but in both cases, replication complexes (red) co-localised precisely with both OAS3 and ZAP (to produce yellow dots) apart from in the immediate perinuclear region. Similarly, although localisations were more diffuse and expression levels different, ZAP co-localised with OAS3. Different combinations of ZAP, OAS3 and dsRNA were co-stained with G3BP (Fig. 9C): in particular the amount of dsRNA co-localised with G3BP appears to be much higher for the CpG-H transfected A549 cells compared to WT transfected cells, higher proposition of dsRNA did not associate with G3BP (to produce distinct red and green punctae)This was confirmed by quantitative analysis (Fig. 9D): for CpG-H and WT replicon transfected A549 cells the co-localisation coefficients were 0.70 and 0.44, respectively. Similarly, substantially less co-localisation of OAS3 with E7 replicon dsRNA was observed in WT E7 transfected cells.

**FIGURE 9.**
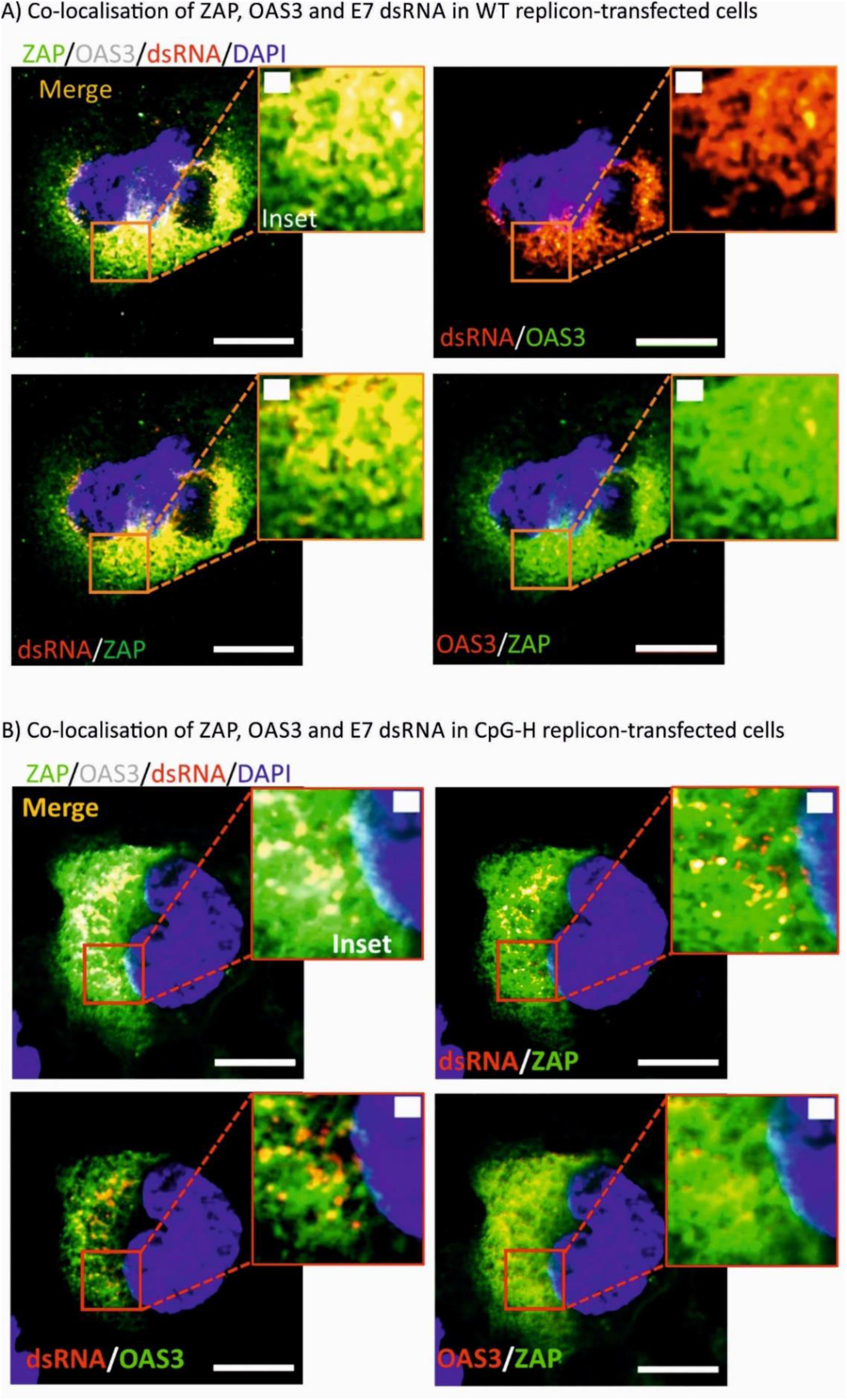
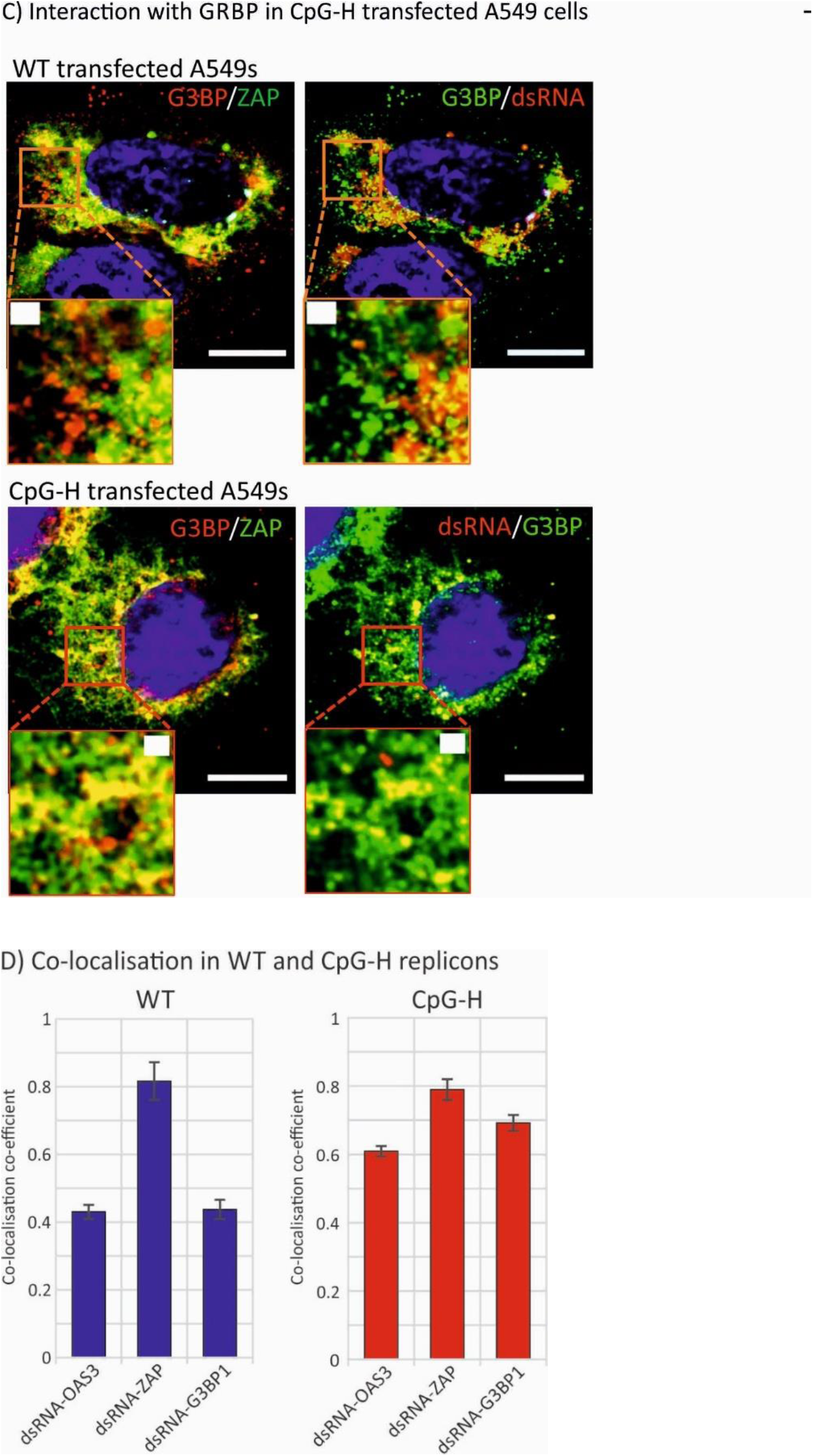
CELLULAR LOCALISATION OF ZAP, OAS3 AND STRESS GRANULE-ASSOCIATED G3BP PRTOEIN. A549 cells were transfected with the (A) WT or (B) CpG-H replicon transcripts and at 6 hours p.t., cells were fixed with 4% PFA and co-stained with ZAP (16820-1-AP, Proteintech), OAS3 (ab188111, Abcam), and dsRNA (J2, SCICONS). Nuclear DNA was stained by DAPI. The scale bars are 10μm and 1 μm, respectively. (C) Visualisation of E7 dsRNA and ZAP localisation with stress granules stained by G3BP (ab220524, Abcam). (D) Quantitation of co-localisation of E7 replicon (dsRNA), ZAP and OAS3, and with G3BP in WT and CpG-H replicon-transfected A549 cells. Bar heights represent the mean of three separate cells fields; error bars show SDs.

## DISCUSSION

### The E7 replicon model

This study used a wide variety of synthetic sequences to characterise the interaction of ZAP and associated cellular proteins to viral RNA and its association with virus attenuation. The use of a previously developed replicon enables effects of cellular restriction of replication through modification of the untranslated 3’UTR to be uncoupled from potential compounding effects of changes in translation efficiency from altered codon usage and codon pair frequencies that might occur in compositionally modified sequences. It also enable much more extensive mutagenesis, since compositional changes do not have to be synonymous and preserve protein coding. The ability to fix G+C content in almost all mutants generated for the project (Table 1) removed a further potential confounding factor influencing replication. The attenuation achieved though modification of the 3’UTR sequence was more restricted (approximately 20-fold for a replicon with an inserted CpG-H sequence; 4-fold for UpA-H) compared to low MOI replication assays with E7 virus. However, the replicon format provides a much more direct, quantitative and better normalised metric of replication ability. Investigation of the effects of compositional modification of the E7 replicon (and the virus) were additionally not compounded by potential unintended effects of sequence modifications on splicing sites and other RNA signal motifs that may be deleted or created *de novo* and compromise protein expression (6).

### Alternative dinucleotides and CpG context and attenuation / ZAP binding

We investigated several questions regarding dinucleotide recognition and virus attenuation, for example, the possibility that attenuation might have arisen through a greater frequency of self-complementary dinucleotides in the sequence and a resulting greater degree of RNA folding as predicted by RNAFold (Table S3; Suppl. Data). The observation of no attenuation and WT-levels of binding to ZAP of mutants enriched for GpC and UpA (Fig. 1B) provides evidence against this possibility. Furthermore, the lack of attenuation and ZAP binding of CpA-H and UpG-H mutants indicates that ZAP-binding, and its apparent affinity for both CpG and UpA dinucleotides, was not simply because it has a broader binding specificity for any pyrimidine followed by any purine. The data showing that UpA and CpG did not compete for binding sites in ZAP (Fig. 6) and that CpG and UpA were not dependent on each other to achieve attenuation (Fig. 7) supports this conclusion.

Recent studies have demonstrated that effects of additional CpG sites on attenuation of HIV-1 was dependent on genome location and positioning within genomic regions forming RNA secondary structures (6, 7). On a smaller scale, we found additionally that attenuation of replicon replication and binding to ZAP mediated through CpG dinucleotides was dependent on their immediate sequence contexts (Figs. 2-4), which the identities of 5’ and 3’ bases exerted a robust effect on replication attenuation and differences in affinity for ZAP binding in the IP assay. These findings suggest that there may be specifically favoured bases around the CpG binding ZAP that promotes binding. It is indeed intriguing, although not commented on in the studies, that both currently published structures of RNA bound to ZAP have U residues 5’ to CpG (23) or at both 5’ and 3’U sites (22), perhaps selected from other candidates because of their propensity to co-crystallise with ZAP.

From analysis of the topology and size constraints of the CpG binding site and immediate upstream and downstream bases (22, 23) It was concluded that there should be no base constraints on the immediate 5’ (−1) base to CpG and not to be part of the CpG binding site (23). Similarly, the topology of both 5’ (−1) are 3’ (+1) bases were considered to not impose any base constraints. Both conclusions fail to match the experimental findings of the current study. It was further predicted that the identity of −2 base (G) was critical for recognition (22). The ZAP structure study was published after our experimental studies were concluded, although there was, in retrospect, no association between ZAP binding affinity (or replication attenuation) with numbers of GNCG motifs in the various constructs analysed in the current study (data not shown). The flexibility of the replicon design will, however, allow these structure-based bindings predictions to be more systematically investigated in the future.

### The ZAP interactome

The study provides a convincing although incomplete explanation for the previously observed co-dependence of ZAP and OAS3 in mediating the attenuation of CpG-H (and UpA-H) mutants of E7. OAS3, while seemingly necessary for attenuation, seems unlikely to be able to bind and discriminate between ssRNA sequences of different dinucleotide compositions, primarily because of its binding specificity to long dsRNA sequences (21). However, an artificial CpG-H (or UpA-H) sequence possess elevated frequencies of self-complementary bases that make the formation of duplexed, internally base-paired stretches of RNA more likely to occur than in native sequences. However, we found that sequences comparably enriched for GpC and ApU sequences with their equally over-represented frequencies of self-complementary bases showed no attenuating effect (Fig. 1B) nor any greater than WT levels of binding to ZAP (Fig. 4).

Indeed, the findings in the current study provided evidence for a cooperative interaction between the two proteins, through their close physical association through their co-precipitation (Fig. 8D), the dependence on ZAP for RNA binding in the OAS3 binding assay (Fig. 8B) and co-localisation of ZAP and OAS3 (and replicon dsRNA) in infected cells (Fig. 9). ZAP has indeed been shown to co-localise with SGs and other cytoplasmic structures (15, 16, 29, 30) and shows a dependence on other proteins, such as TRIM25 and the nuclease, KHNYN for its antiviral activity against HIV-1 (6, 8). However, the 78 cellular proteins, many associated with stress granules, that were verified to co-IP with ZAP on mass spectrometry analysis (15) indicates its much larger interactome and intimate connection with other antiviral, stress response and RNA degradation pathways. Amongst the latter, co-IP and in some cases direct binding with ZAP were observed for the 3’-52 exosome component EXOSC8 (RRP43), EXOSC5 (RRP46), the helicase DHX30 and the 5’-3’ exoribonuclease 2 (XRN2). While TRIM25 was identified in that list, KHNYN and OAS3 were not, despite the clear evidence for their co-precipitation in targeted WB assays ((6); Fig. 8D). In view of the latter findings, perhaps this analysis might be re-visited.

The study provides further evidence for the attenuating effects of elevated frequencies of the UpA dinucleotide on E7 replication and binding in ZAP and OAS3 IP assays (Figs. 1B, 3, 6, 7). These observations contrast with predictions from recent structural studies that UpA could not be accommodated within the ZAP binding site as an alternative to CpG (22, 23). The possible existence of an entirely separate UpA binding site on ZAP or associated protein was indeed experimentally supported by the observed absence of binding of alternative YpR dinucleotides (Fig. 1B) and the lack of competition by CpG- and UpA-enriched RNA sequences for binding sites in competition assays (Fig. 6). It is conceivable that the complex of ZAP-associated proteins identified by IP may include one or more further RNA binding proteins that could mediate the apparently specific recognition of UpA-enriched sequences, including those with zinc finger domains in the ZAP interactome (15) such as zinc finger RNA-binding protein (31). However, the possible existence of an alternative UpA recognition protein in the larger stress granule / ZAP complex does not square with the apparent dependence on ZAP expression in the OAS3 IP assay (Fig. 8B). In the ZAP k/o (B8) cell line, only WT background levels of binding were observed for the UpA-H (and CpG-H) transcript, that contrasts the 2 log greater binding in ZAP-expressing A549 cells (Fig. 8A).

There are undoubtedly vast complexities in the cellular response to RNA sequences of different compositions and configurations. For example, ZAP binding may directly or indirectly activate OAS3 independently of RNAseL; this might represent a complementary or additional effector pathway to KHNYN, perhaps targeting RNA in the different cellular compartments of E7 in replication complexes and HIV-1 sequences expressed as mRNAs. The use of a range of different viral models with their different replication strategies is clearly essential to fully understand the nature and purpose of CpG- and UpA-mediated restriction of virus replication in mammalian cells.

## METHODS

### Cells, viruses and reagents

Cells were maintained in Dulbecco’s modified Eagle medium (DMEM) (Life Technologies) supplemented with 10% heat-inactivated foetal calf serum (FCS) (Gibco), 100 U/ml penicillin and 100 µg/ml streptomycin (Life Technologies). A549 ZAP knockout (k/o) and A549 OAS3 k/o cell lines with deletion of all three for all three alleles (-/-/-) were grown as described previously (1).

### Virus and replicon assays

The design and construction of E7 isolate Wallace infectious clone and replicon with various nucleotide compositions in R1 was described previously (20, 32). All R1 insert sequences used in the study are provided in FASTA format (Table. S1; Suppl. Data). RNA preparation and assessment of replication ability were performed as previously described {#30969}. Quantitative real-time PCR (qRT-PCR), SDS-PAGE and immunoblotting were performed as previously described {#30969}.

### Immunofluorescence (IF)

IF assays were performed as previously described {#30969}, using cells seeded onto 19 mm glass coverslips in 24 well plates., 6 h post transfection cells were fixed in 4% PFA and permeabilised with 0.1% (v/v) Triton X-100 (Sigma-Aldrich)in PBS for 15 min. Coverslips were washed in PBS and blocked in blocking buffer (3% (w/v) BSA in PBS) and the primary antibody applied at the relevant dilution in 1% (w/v) BSA (Sigma) in PBS and incubated overnight at 4 °C. To remove any unbound primary antibody, cells were washed three times in PBS before the application of the relevant Alexa Fluor-488, 594 or 647 conjugated secondary antibodies (Life Technology) diluted 1:750 in 1% (w/v) BSA in PBS followed by 2 h incubation at RT in the dark. DAPI staining and microscopy were performed as previously described {#30969}.

### RNA-binding protein immunoprecipitation (IP)

IP **in the previously used intracellular format** were performed as previously described {#30969}. For *ex vivo* assay format, cell lysates were first prepared from monolayer cultures with or without **interferon (**IFN)-β (eBioscience) stimulation (100 ng/mL) 24 hours before harvesting in 15 cm dishes using RIP lysis buffer containing protease and RNAse inhibitors (Millipore). Cell lysates and RNA transcripts (at indicated concentration) mixture for each reaction was then incubated with magnetic beads A/G associated either with rabbit anti-ZAP (5 µg per/reaction; Proteintech) or normal rabbit IgG (Millipore) as described above.

For the competitive RNA binding assay, four binding assays were conducted for each experiment. Cell lysates were incubated with CpG-H E7 transcripts at 1:0.1, 1:1, 1:10, and 1:100 molar mass ratios to UpA-H and *vice versa* (UpA-H E7 transcripts at indicated molar mass ratios to CpG-H). Cell lysates were incubated for 5 h and the samples were washed six times, with complete resuspension of beads between washes and 15 min incubation with rotation for the last two washes. After washings, the samples were re-suspended in RIP wash buffer and divided in aliquots. A 100 µl aliquot of immunecomplexes was reserved for analysis of the protein fraction. After beads were immobilized, the wash media were discarded, and beads were re-suspended in Laemmli sample buffer. About 10 µl of the inputs were also mixed with Laemmli buffer. All the samples were heated for 5 min at 95°C and separated by SDS-PAGE for western blot analysis. In parallel, viral RNA were purified from another aliquot of immune-complexes and inputs, using the RNAviral kit (Zymo research). RNAs were eluted in 15 µl of RNase–DNase free water. The copy number of E7 present in each fraction was determined by qPCR as described above.

## ACKNOWLEDGEMENTS

We thank Harriet Thomas and Jack White for technical assistance. We are grateful to Paul Klenerman group from the Nuffield Department of Medicine for advice, for access to the facility/equipment, and providing IFN-β.

## SUPPLEMENTARY DATA CONTENTS

TABLE S1. Composition of synthetic ncR1 insert sequences

TABLE S2. Primers used for E7 RNA quantitation

FIGURE S1. Replication abilities of YpR and reverse dinucleotide mutants in RD cells

FIGURE S2. Replication abilities of CpG context mutants in RD cells

FIGURE S3. Specificity of amplification of CpG-H and UpA-H R1 RNA sequences

## REFERENCES

1. Odon V, Fros JJ, Goonawardane N, Dietrich I, Ibrahim A, Alshaikhahmed K, Nguyen D, Simmonds P. 2019. The role of ZAP and OAS3/RNAseL pathways in the attenuation of an RNA virus with elevated frequencies of CpG and UpA dinucleotides. Nucleic Acids Res 47:8061–8083.

2. Trus IU, D.; Berube, N.; Cheler, C.; Martel, M..-J.; Gerdts, V.; Karnychul, U. 2020. CpG-Recoding in Zika Virus Genome Causes Host-Age-Dependent Attenuation of Infection With Protection Against Lethal Heterologous Challenge in Mice. Front Immunol https://doi.org/10.3389/fimmu.2019.03077.

3. Kozaki T, Takahama M, Misawa T, Matsuura Y, Akira S, Saitoh T. 2015. Role of zinc-finger anti-viral protein in host defense against Sindbis virus. Int Immunol 27:357–64.

4. Bick MJ, Carroll JW, Gao G, Goff SP, Rice CM, MacDonald MR. 2003. Expression of the zinc-finger antiviral protein inhibits alphavirus replication. J Virol 77:11555–62.

5. Chiu HP, Chiu H, Yang CF, Lee YL, Chiu FL, Kuo HC, Lin RJ, Lin YL. 2018. Inhibition of Japanese encephalitis virus infection by the host zinc-finger antiviral protein. PLoS Pathog 14:e1007166.

6. Ficarelli M, Wilson H, Pedro Galao R, Mazzon M, Antzin-Anduetza I, Marsh M, Neil SJ, Swanson CM. 2019. KHNYN is essential for the zinc finger antiviral protein (ZAP) to restrict HIV-1 containing clustered CpG dinucleotides. Elife 8.

7. Kmiec D, Nchioua R, Sherrill-Mix S, Sturzel CM, Heusinger E, Braun E, Gondim MVP, Hotter D, Sparrer KMJ, Hahn BH, Sauter D, Kirchhoff F. 2020. CpG Frequency in the 5’ Third of the env Gene Determines Sensitivity of Primary HIV-1 Strains to the Zinc-Finger Antiviral Protein. mBio 11.

8. Takata MA, Goncalves-Carneiro D, Zang TM, Soll SJ, York A, Blanco-Melo D, Bieniasz PD. 2017. CG dinucleotide suppression enables antiviral defence targeting non-self RNA. Nature 550:124–127.

9. Miyazato P, Matsuo M, Tan BJY, Tokunaga M, Katsuya H, Islam S, Ito J, Murakawa Y, Satou Y. 2019. HTLV-1 contains a high CG dinucleotide content and is susceptible to the host antiviral protein ZAP. Retrovirology 16:38.

10. Mao R, Nie H, Cai D, Zhang J, Liu H, Yan R, Cuconati A, Block TM, Guo JT, Guo H. 2013. Inhibition of hepatitis B virus replication by the host zinc finger antiviral protein. PLoS Pathog 9:e1003494.

11. Zhu Y, Wang X, Goff SP, Gao G. 2012. Translational repression precedes and is required for ZAP-mediated mRNA decay. Embo j 31:4236–46.

12. Guo X, Ma J, Sun J, Gao G. 2007. The zinc-finger antiviral protein recruits the RNA processing exosome to degrade the target mRNA. Proc Natl Acad Sci U S A 104:151–6.

13. Zhu Y, Chen G, Lv F, Wang X, Ji X, Xu Y, Sun J, Wu L, Zheng YT, Gao G. 2011. Zinc-finger antiviral protein inhibits HIV-1 infection by selectively targeting multiply spliced viral mRNAs for degradation. Proc Natl Acad Sci U S A 108:15834–9.

14. Moldovan JB, Moran JV. 2015. The Zinc-Finger Antiviral Protein ZAP Inhibits LINE and Alu Retrotransposition. PLoS Genet 11:e1005121.

15. Goodier JL, Pereira GC, Cheung LE, Rose RJ, Kazazian HH, Jr. 2015. The Broad-Spectrum Antiviral Protein ZAP Restricts Human Retrotransposition. PLoS Genet 11:e1005252.

16. Lee H, Komano J, Saitoh Y, Yamaoka S, Kozaki T, Misawa T, Takahama M, Satoh T, Takeuchi O, Yamamoto N, Matsuura Y, Saitoh T, Akira S. 2013. Zinc-finger antiviral protein mediates retinoic acid inducible gene I-like receptor-independent antiviral response to murine leukemia virus. Proc Natl Acad Sci U S A 110:12379–84.

17. Zheng X, Wang X, Tu F, Wang Q, Fan Z, Gao G. 2017. TRIM25 Is Required for the Antiviral Activity of Zinc Finger Antiviral Protein. J Virol 91.

18. Sanchez JG, Sparrer KMJ, Chiang C, Reis RA, Chiang JJ, Zurenski MA, Wan Y, Gack MU, Pornillos O. 2018. TRIM25 Binds RNA to Modulate Cellular Anti-viral Defense. J Mol Biol doi:10.1016/j.jmb.2018.10.003.

19. Li MM, Lau Z, Cheung P, Aguilar EG, Schneider WM, Bozzacco L, Molina H, Buehler E, Takaoka A, Rice CM, Felsenfeld DP, MacDonald MR. 2017. TRIM25 Enhances the Antiviral Action of Zinc-Finger Antiviral Protein (ZAP). PLoS Pathog 13:e1006145.

20. Fros JJ, Dietrich I, Alshaikhahmed K, Passchier TC, Evans DJ, Simmonds P. 2017. CpG and UpA dinucleotides in both coding and non-coding regions of echovirus 7 inhibit replication initiation post-entry. Elife 6.

21. Donovan J, Whitney G, Rath S, Korennykh A. 2015. Structural mechanism of sensing long dsRNA via a noncatalytic domain in human oligoadenylate synthetase 3. Proc Natl Acad Sci U S A 112:3949–54.

22. Luo X, Wang X, Gao Y, Zhu J, Liu S, Gao G, Gao P. 2020. Molecular Mechanism of RNA Recognition by Zinc-Finger Antiviral Protein. Cell Rep 30:46-52.e4.

23. Meagher JL, Takata M, Goncalves-Carneiro D, Keane SC, Rebendenne A, Ong H, Orr VK, MacDonald MR, Stuckey JA, Bieniasz PD, Smith JL. 2019. Structure of the zinc-finger antiviral 1. protein in complex with RNA reveals a mechanism for selective targeting of CG-rich viral sequences. Proc Natl Acad Sci U S A 116:24303–24309.

24. Coleman JR, Papamichail D, Skiena S, Futcher B, Wimmer E, Mueller S. 2008. Virus attenuation by genome-scale changes in codon pair bias. Science 320:1784–7.

25. Mueller S, Coleman JR, Papamichail D, Ward CB, Nimnual A, Futcher B, Skiena S, Wimmer E. 2010. Live attenuated influenza virus vaccines by computer-aided rational design. Nat Biotechnol 28:723–6.

26. Ni YY, Zhao Z, Opriessnig T, Subramaniam S, Zhou L, Cao D, Cao Q, Yang H, Meng XJ. 2014. Computer-aided codon-pairs deoptimization of the major envelope GP5 gene attenuates porcine reproductive and respiratory syndrome virus. Virology 450-451:132–9.

27. Martrus G, Nevot M, Andres C, Clotet B, Martinez MA. 2013. Changes in codon-pair bias of human immunodeficiency virus type 1 have profound effects on virus replication in cell culture. Retrovirology 10:78.

28. Le Nouen C, Brock LG, Luongo C, McCarty T, Yang L, Mehedi M, Wimmer E, Mueller S, Collins PL, Buchholz UJ, DiNapoli JM. 2014. Attenuation of human respiratory syncytial virus by genome-scale codon-pair deoptimization. Proc Natl Acad Sci U S A 111:13169–74.

29. Chen S, Xu Y, Zhang K, Wang X, Sun J, Gao G, Liu Y. 2012. Structure of N-terminal domain of ZAP indicates how a zinc-finger protein recognizes complex RNA. Nat Struct Mol Biol 19:430–5.

30. Todorova T, Bock FJ, Chang P. 2014. PARP13 regulates cellular mRNA post-transcriptionally and functions as a pro-apoptotic factor by destabilizing TRAILR4 transcript. Nat Commun 5:5362.

31. Haque N, Ouda R, Chen C, Ozato K, Hogg JR. 2018. ZFR coordinates crosstalk between RNA decay and transcription in innate immunity. Nat Commun 9:1145.

32. Atkinson NJ, Witteveldt J, Evans DJ, Simmonds P. 2014. The influence of CpG and UpA dinucleotide frequencies on RNA virus replication and characterization of the innate cellular pathways underlying virus attenuation and enhanced replication. Nucleic Acids Res 42:4527–45.

